# Active manipulation of surface orientation under directional illumination improves color discrimination

**DOI:** 10.64898/2026.11.03.763965

**Authors:** Sheikh Mohiddin, Takuma Morimoto, Fabian Coupette, Hannah E. Smithson

**Affiliations:** University of Oxford, Department of Experimental Psychology, Oxford, UK

## Abstract

Two surfaces that appear different colors under one illumination might appear the same color under another illumination. This phenomenon is known as illuminant metamerism and is thought to limit surface color discrimination. However, real-world lighting environments are not directionally uniform. In theory, surfaces that appear the same color at one tilt might not appear the same color at another tilt where they sample a different illumination spectrum. We tested whether human observers could use this directional variation to improve surface color discrimination. Our results confirm that human color discrimination is improved when observers are able to tilt surfaces to sample directional lighting. Observers’ sampling motions were spatiotemporally structured and appeared to be directed towards the general lighting geometry rather than systematically maximizing color discriminating information for specific surface pairs.

## 1. Introduction

Our ability to distinguish objects by their spectrally selective reflectance is limited by our biology. Color perception depends on the responses of wavelength-sensitive cone cells in the human retina. With three classes of cone cells, an incoming spectrum of light is described by only a triplet of signals. As the univariant cone responses confound the wavelength and intensity of incoming light [1], two lights that differ in spectral composition might nevertheless elicit the same triplet of cone signals and thus appear the same color. The same principle extends to the perception of surface color, leading to a phenomenon called *illuminant metamerism*: two surfaces with distinct reflectance spectra might elicit identical cone responses under one illuminant, but distinct cone responses under another. A notable example of this is when choosing matching textiles: a customer might choose items of clothing whose colors match under indoor store lighting, but look distinct under daylight.

A surface that appears a certain color under one illuminant could appear a range of other colors under other illuminants. Theoretical estimates of the frequency and extent of illuminant metamerism suggest that it could significantly impair the ability to discriminate surfaces by their reflectance spectra and challenge the notion of constant color perception [2]. However, when considering illuminants and surface reflectances that are commonly encountered in natural environments, illuminant metamerism appears to be less of a hindrance [3, 4].

Many characterizations of illuminant metamerism consider single, uniform illuminations. Yet, lighting in realistic environments exhibits directional spectral variability—surfaces receive different spectral illumination depending on their surface tilt [5]. Whereas a pair of surfaces might elicit identical cone signals at one *tilt*, their cone signals could diverge at *another tilt* (Fig. 1). Recent simulations show that the spectral variation present in some natural environments is sufficient to break most cases of metamerism; pairs of surfaces that are indistinguishable at one tilt will typically be distinguishable at another tilt [6]. Directional illumination provides information that, in theory, could be practically accessed by simply tilting surfaces [7].

**Fig. 1.**
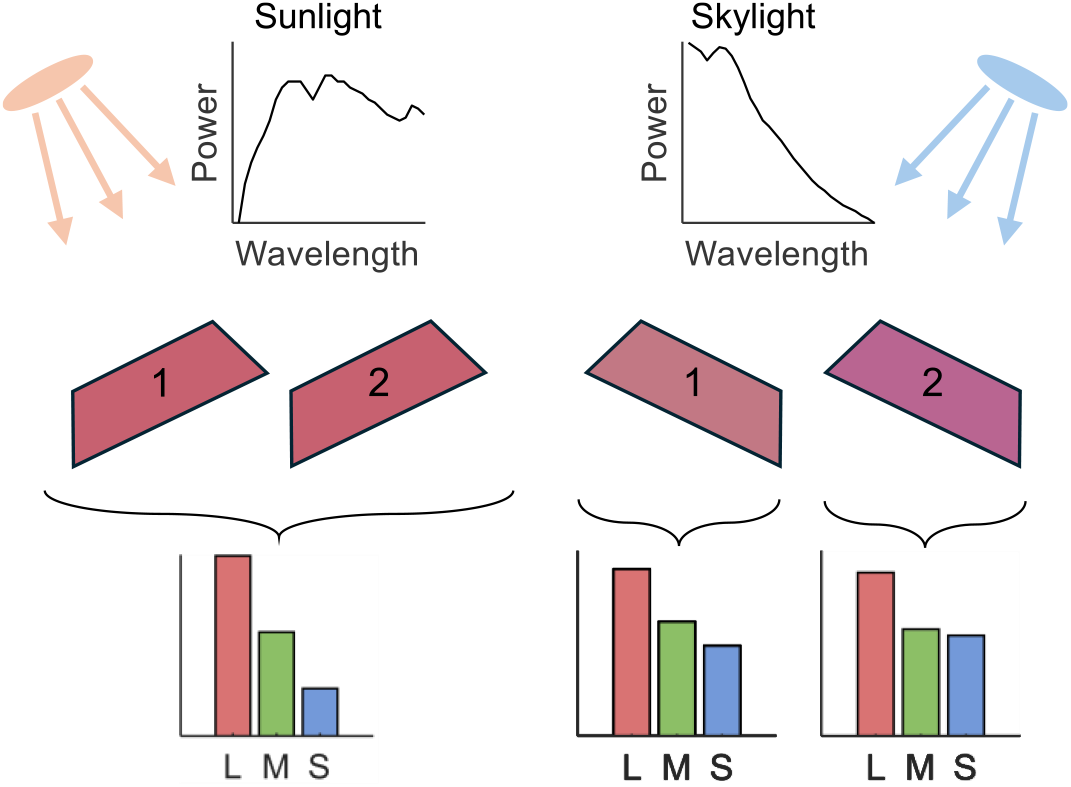
Two surfaces with different reflectance (1 and 2) functions tilted towards sunlight might elicit the same cone signals and appear identical (left). When tilted towards diffuse skylight (right), the reflected light elicits different cone signals, increasing the surfaces’ discriminability.

Observing multiple views of a stimulus generally supports stable perception. From a sequence of retinal images, each corresponding to a transformation of a stimulus, an observer can infer which aspects of the presentation are due to stable object properties, and which aspects are due to the environment [8,9]. Studies into color perception have typically characterized responses to static stimuli, while those employing dynamic stimuli typically present pre-determined, experimenter-controlled temporal transformations of stimuli, such as changes in illuminant, chromatic modulation, or feature synchrony [3,10–12]. In contrast, observers readily employ *active* sensing: actively and intentionally sampling new information to make more accurate estimates about the properties of objects and the environment, thereby resolving perceptual ambiguities. This approach has been broadly demonstrated across fields such as material perception [13–15], eye movements [16–18], as well as reinforcement learning [19]. An additional question in such studies is what strategies, if any, observers use to gather information.

Relatively little is known about color perception when observers can actively manipulate the visual information they receive. Could people actively exploit directional illuminant variation? This study tested color discrimination of physical samples in an environment with directionally varying illumination. Observers performed a color discrimination task under different viewing conditions: either they could freely tilt surfaces, or they observed surfaces at fixed tilts. Tilting trajectories were quantitatively measured to characterize task-relevant strategies. Using physically based simulation techniques, the tilting trajectories were also analyzed with respect to the available color difference signals at each timepoint. The experiment tested the hypotheses that people actively sought tilts where the surfaces were more discriminable, and that finding these tilts would improve performance.

## 2. Methods

### 2.1 Participants

The study was approved by the Central University Research Ethics Committee at the University of Oxford (RE77333/RE009). Each participant received a full explanation of the experimental procedure and provided informed consent prior to the experiment. Twenty-four participants (four males) between 19 and 34 years of age (M = 22.2 yrs, SD = 4.0) with normal or corrected-to-normal visual acuity were recruited for the experiment. Color vision was tested using Hardy-Rand-Rittler (HRR) pseudoisochromatic test plates [20] under a D65 illuminant. Four participants’ datasets were removed due to experimenter error, poor visual acuity, voluntary drop-out, and failing to follow the task instructions. Data from twenty participants (three males) between 19 and 34 years of age (M = 22.3 yrs, SD = 4.0) were retained for analysis.

### 2.2 Apparatus

Participants performed a three-alternative forced choice task (3AFC), i.e. an odd-one-out task. The participants viewed the stimuli in an open-front-face viewing box in a dark room. The viewing box had a 50 × 50 cm base, and a height of 60 cm. Two LED bulbs (LIFX Color 1100 lumens, model LHLA19E26US, LIFX, Pico Rivera, CA) were mounted 13cm from the left and right edges of the top of the box (Fig. 2a). The lights were set to correlated color temperature of 1500 K and 9000 K (1500 K *xy*-chromaticity: 0.592, 0.381; 9000 K: 0.275, 0.285) and matched for luminance (161.3 cd/m^2^ for the 1500 K light at full brightness and 9000 K light at 46.2% brightness). This arrangement provides the directional illuminant variability within the box, and mimics an exaggerated warm–cool axis that might be observed when tilting between warm direct sunlight and cool diffuse skylight. A layer of Clairefontaine Maya black paper was placed over a portion of the open front face to occlude the light sources, and the same paper was used to cover the inner walls of the box to reduce interreflection.

**Figure 2.**
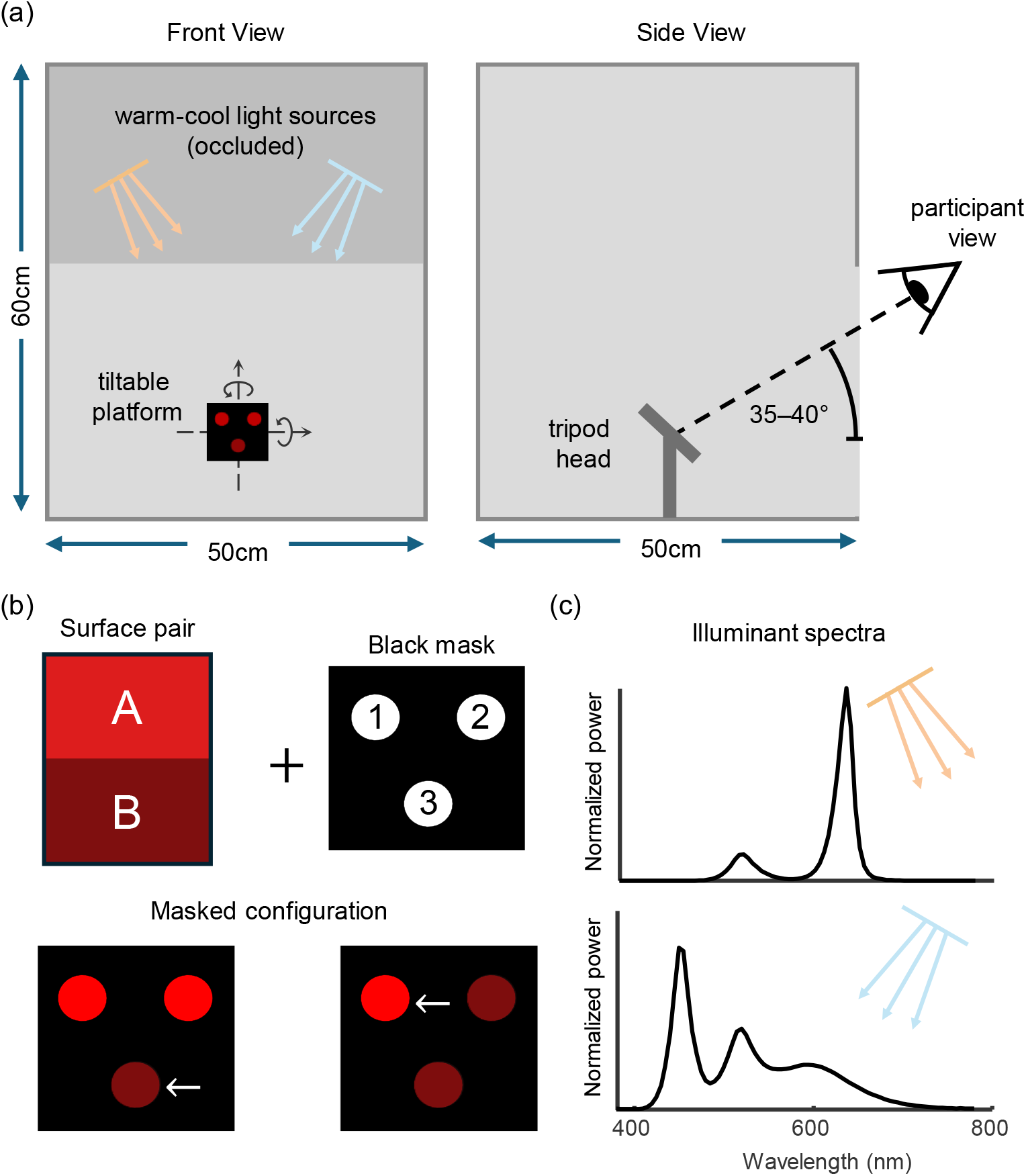
(a) Schematic of the apparatus. (b) Each surface pair can be configured six ways by choosing which surface is the odd one out (white arrow) and where it is located. (c) Illuminant spectra for the warm and cool LEDs.

A tripod ball head (K&F CONCEPT B234A1+BH-28, Shenzhen Zhuoer Photography Equipment Co., Ltd., Shenzhen, China) was fixed to the center of viewing box base. This tripod head allowed tilts of up to 40° from vertical, excluding a drop notch positioned away from the participant. A 10cm square, black paper-covered platform was attached to the tripod head. Participants were seated directly in front of the tripod with their head on a chinrest. The height of the chinrest and distance of the viewing box were adjusted until participants could comfortably move the platform through its whole range of motion.

Participants viewed three circular color samples, each subtending a visual angle of 0.7 degrees (larger if participants moved closer). An 8 × 8 cm cardboard platform was attached magnetically to the larger moving platform. This mask had 8 mm circular cutouts whose centers were 1.7 cm from each other, and 1cm from the platform center. A pair of adjoined paper paint swatches were attached magnetically to the back of the mask, such that one material would appear in two positions, and the other material would appear in the third position. The pair could be rotated to one of six possible configurations in the plane of the platform, changing the identity (surface *A* or surface *B*) and position (one out of three) of the distinct material. The surface of the mask was covered with black paper, except for the 8 mm circular cutouts, and the back of the cardboard platform was cut out so that the colored surfaces were near flush with the black paper (Fig. 2b).

Trajectories were measured using a development board (XIAO nRF52840 Sense, Seeed Studio, Shenzhen, China) housing a 6-axis inertial measurement unit (IMU) sampling at 62.5 Hz. The board was mounted behind the far-right corner of the platform and connected to the experimenter’s laptop using a matte black USB-C cable. The cable was routed to avoid interfering with the platform’s motion. The experimenter’s laptop screen was set to the lowest brightness and positioned to minimize light reflecting back into the box.

### 2.3 Stimulus appearance simulation

Stimuli were chosen from 132 different paper paint swatches from the Farrow & Ball Signature Palette. Measurements were taken using a spectroradiometer (PR-670, Photo Research Inc., Chatsworth, CA), under a D65 lamp in a light booth (Judge QC, X-Rite, Grand Rapids, MI), and the illuminant spectrum was measured via a diffuse white standard (Spectralon ®, Labsphere, North Sutton, NH). The spectroradiometer and white standard were also used to measure illuminant spectra for the 1500 K and 9000 K LEDs (Fig. 2c).

The purpose of the simulation was to estimate the color of every patch at every tilt that participants could reach, so that the proximal color-difference signal—the distinctiveness of the target patch (Fig. 2b)—available to the observer could be computed for any platform orientation recorded during the experiment. The subsequent computer graphics simulation techniques were based on those used in a previous study [6]. The physical scene was recreated to scale in the modeling software Blender (version 4.3.2; Blender Foundation, 2024), and only included the padded walls of the box, the lights, and the color patches. All surfaces were modeled as perfectly diffuse, and the lights were modeled as area lights (Fig. 3a).

**Fig. 3.**
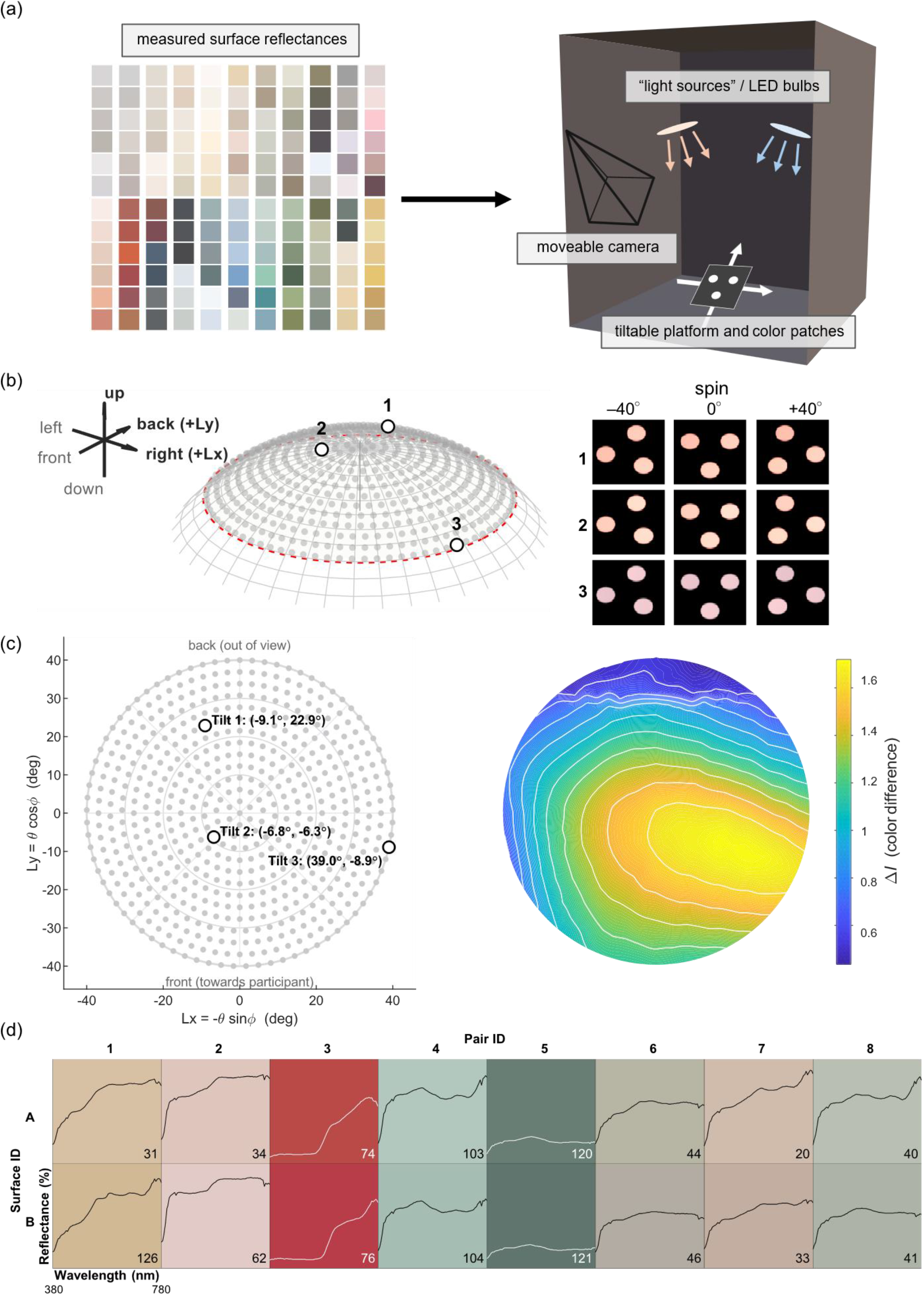
(a) The physical scene was created in 3D modeling software, and the optical interactions were simulated hyperspectrally (b) Left: the sampled 562 tilts are represented as points on surface of a 40° spherical cap (red dashed line). Right: The three rows refer to the three example tilts indicated as 1, 2 and 3 on the spherical cap. The central column, labelled spin 0°, shows the three angular positions of the three patches at the start of a trial. The left and right columns correspond to the spin of the platform by –40° and +40°, respectively. Collectively, these are the 9 rendered stimulus positions at each tilt. (c) The projection (logarithmic map) from the spherical surface to a 2D plane, and a visualization of the color difference at each tilt. The contour lines are lines of constant color difference. (d) The final selection of surface pairs. Their reflectance functions over the wavelength range from 380 nm to 780 nm are overlaid. Inset: their number in the Farrow & Ball Signature Palette (column-major order).

The color patches were transformed into three degrees of rotational freedom: spin around the platform’s *z*-axis, sampled at nine angular positions 1cm away from the platform center, and 562 tilt positions within the 40° spherical cap (Fig. 3b). These tilt positions were chosen to be roughly 3° apart: we first chose 40/3° ≈ 13 concentric rings around the pole and then chose 120 ⋅ sin *θ* ° points for the ring at *θ*°. The nine angular spin positions allow the appearance to be simulated for different distractor/patch target locations, as well as accounting for any incidental platform spin (Fig. 3b). The scene camera tracks the tilting surfaces and renders their appearance at a fixed offset to maximize pixel counts and reduce render noise. Because the surfaces were modelled as perfectly diffuse (Lambertian), their apparent color depended only on the illumination geometry and was independent of the observer’s position.

The patches were rendered hyperspectrally [21] and spectral outputs were converted to CIE 1931 images [22]. For each configuration, the mean XYZ of the color patch was calculated and converted to CIELAB space [23]. The white point values were based on the average tristimulus values of a rendered perfect white diffuse surface at the vertical rest pose. In total, we simulated L*a*b* values for 132 colored surfaces at 562 tilts and 9 angular positions.

### 2.4 Color difference calculation

Colorimetric calculations were performed in MATLAB version 2024b [24] using Psychtoolbox version 3.0.19 [25]. To quantify the available information for a given trial configuration with three visible patches, *i, j*, and *k*, we adapted the CIEDE2000 formula [26] into a three-way difference measure. Surface *i* is assigned a difference measure that is the mean of its color difference Δ*E*^*^ with the other two surfaces *j* and *k*, which specifies how distinct a given patch is from the other two patches.

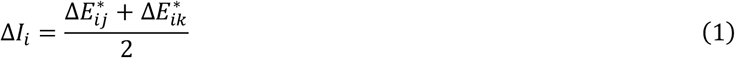

The overall difference measure per trial configuration is based on a combination of the three-way measure corresponding to each patch (C: correct patch; D1, D2: distractor patches).

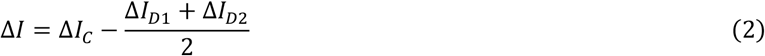

This measure quantifies the extent to which the correct patch is the most distinct; intuitively, if the three patches are similarly distinct, Δ*I* tends towards zero whereas, if the correct patch is more distinct, Δ*I* increases. Note that although the distractor patches’ reflectance functions are identical, they may not always reflect the same light in the physical setup.

For each possible stimulus configuration, the Δ*I* value can be calculated and across the full range of transformations. We chose thresholds of Δ*I* = 1.25, 1.50 and 1.75, and sought to find combinations of surfaces that would be below threshold at most visible tilts, regardless of which surface within the pair was the target, and where the target was located (Fig. 2b). These criteria were intentionally conservative to make trials more difficult and encourage participants to seek out disambiguating color signals. The final selection criteria sought to maximize chromatic variation. The experimenters (the authors) verified and refined the selection to eight surface pairs (Fig. 3d). With two possible odd-one-out identities and three target positions for the odd one out (Fig. 2b), there were 48 possible stimulus configurations.

### 2.5 Experimental design and procedure

Participants performed 48 trials of a three-alternative forced-choice task in which they judged, without feedback, which of three simultaneously presented color patches was the odd one out. The main manipulation was the viewing condition, split into two categories: fixed blocks, in which the participant made a judgment without moving the platform, and free blocks, in which the participant could freely tilt the platform before making a judgment. Fixed blocks were further split into “easy” and “hard” trials, in which the platform’s position corresponded to a relatively high or low Δ*I* for the stimulus configuration. This separation established a baseline that could contextualize the relative strength of a free-viewing effect between two extremes.

Participants performed four blocks of 12 trials in which they saw all 48 possible stimulus configurations. Halfway through the experiment, the lighting arrangement was mirrored (the warm light was switched to cool and vice versa). Participants performed one fixed-viewing block and one free-viewing block under each lighting arrangement. The starting light arrangement, the block order and stimulus order were randomized, avoiding consecutive presentations of the same surface pair.

Before each session, randomized trial tables were generated, including the tilts for the easy and hard trials. The LEDs were set to the starting light arrangement and the main lights in the experiment room were switched off. The IMU was powered on to warm up, and session calibration readings were recorded. The platform was configured with an example color pair separate from the main set of eight.

Participants were seated in front of the viewing box and were instructed to adjust the height of the chinrest and chair for comfort. The viewing box was moved closer to ensure they could move the platform through its whole range of motion. An explanation of the task was given using the example pair. The participant was told that “two of these patches come from the same piece of paper, whereas the other one comes from a different piece of paper”. This phrasing encourages participants to make judgments about the physical properties of the objects, rather than their associated sensations, such as simply which looks different [27]. Participants performed an example fixed trial then an example free trial. During free trials, the platform started away from the participants, out of view, and the participant had to move it into view. Participants were told to freely tilt the platform but to avoid excessive “spinning” movements. Participants were asked to stop moving the platform before indicating their answer.

After the participant had understood how to move the platform comfortably, the main experiment began. Between each trial, participants were asked to close their eyes. During this time, the experimenter configured the next trial based on the color pair, the odd one out position, the odd one out identity, and starting position. Before fixed trials, feedback from the IMU guided the experimenter to the target tilt. If the angle was too extreme, the experimenter asked the participants if all three patches could be seen. If not, the platform was moved to a less extreme angle of similar relative difficulty. IMU recording was started manually before free trials and manually ended once the participant gave their answer. Participants were asked to temporarily leave the room while the light arrangement was switched.

### 2.6 Data analysis

Generalized linear mixed-effects models (GLMMs) were fit on response accuracy, using a binomial distribution with a logit link function. The main model predicted response accuracy as a function of fixed effects for viewing condition and color pair, and a random effect of participant. Color pair was added to account for inherent difficulty differences between certain pairs of colors.

Trajectories were reconstructed using the 6-axis IMU values. The accelerometer measures three axes’ orientations with respect to gravity. Ratios between the measurements describe the platform’s tilt with two degrees of freedom. The gyroscope measures the platform’s angular velocity in each axis. Integrating the gyroscope readings provides the final degree of freedom for describing the platform’s orientation. Session calibration readings provide an estimate for any fixed gyroscope bias in each axis and are subtracted from the raw readings at each timepoint to prevent accumulation of bias while integrating. The “bringing-in” window (the starting out-of-view component and the first 200ms of in-view trajectory) was cropped, along with the period coming more than 150ms after the last sustained stationary tilt.

With three degrees of freedom, the participants’ 3D manipulations can be fully reconstructed. Each transformation of the platform corresponds to a particular Δ*I*, interpolated from the simulated set of 562 tilts and angular positions for the stimulus configuration. Reading out the interpolated Δ*I* produces an overall “Δ*I*-trajectory”: a time series representing the physical distinctiveness of the target patch at each point in the trajectory. For trajectories under flipped lighting arrangement, the stimulus configuration is changed accordingly before reading out Δ*I*: left and right odd-one-out positions are switched, surface spin is negated, and the tilt position is left–right mirrored.

Subsequent motion analyses mainly focused on tilting motions, while spin was only used to improve the estimate of the surface appearance. These tilt trajectories exist on the curved spherical surface (Fig. 3b). We encoded them as (Lx, Ly) coordinates on the logarithmic map of the sphere, which preserves distances and directions from the pole (Fig. 3c). Trajectories were analyzed in three main ways: whether scalar descriptors of the sampled information could predict performance; whether tilting trajectories were aligned with the spatial geometry of available color signals; and whether sampling patterns were spatiotemporally structured.

## 3. Results

The results are organized around two questions. We first ask whether being able to tilt the surfaces improved color discrimination (Sections 3.1 and 3.2). We then ask how observers sampled the available color-difference signals through tilting, from three complementary approaches: whether simple summaries of the sampled Δ*I* predict accuracy (Section 3.3), whether tilting movements were directed toward tilts that increase Δ*I* (Section 3.4), and whether the sampling had a temporal structure (Section 3.5).

### 3.1 Descriptive statistics for viewing condition

We first summarize raw accuracy across viewing conditions. The two fixed-viewing conditions provide reference points: easy-fixed trials indicate performance when the platform is already at an informative tilt, whereas hard-fixed trials indicate performance at an uninformative tilt. We then ask how performance under free viewing compares with these two reference conditions. The final sample comprised 20 participants and 973 trials (492 free-viewing, 240 easy-fixed, 241 hard-fixed; one participant performed an additional 13 trials, which were retained for analysis). Mean accuracy was highest under easy-fixed viewing (M = 0.68, SD = 0.16), similar under free viewing (M = 0.65, SD = 0.15), and lowest under hard-fixed viewing (M = 0.51, SD = 0.14). Participant accuracy across all conditions ranged from 0.25 to 0.96 (Fig. 4a). Between different color pairs, mean accuracy across all conditions ranged from 0.38 to 0.89 (Fig. 4b). In short, when free to tilt the surfaces, observers performed considerably better than at an uninformative fixed tilt and nearly as well as at an informative fixed tilt, although there was substantial variability across participants and across color pairs.

**Figure 4.**
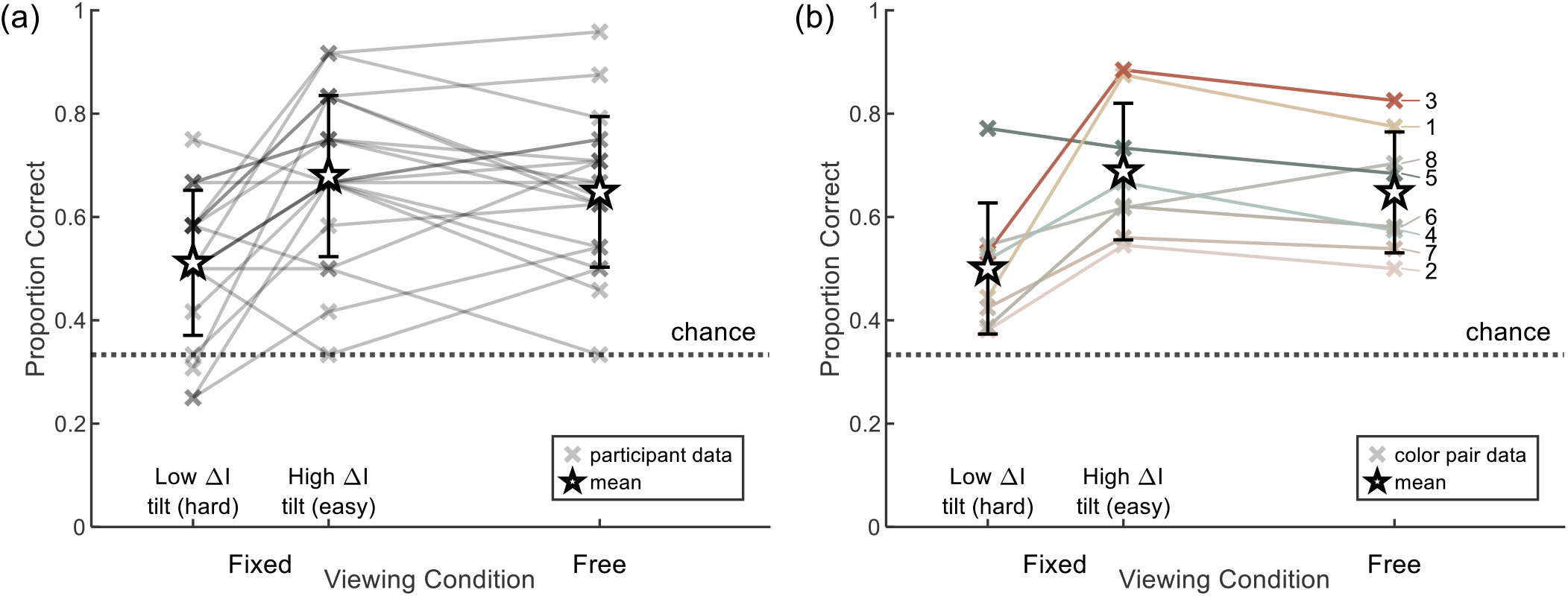
Mean accuracy for each viewing condition. Individual data are shown with crosses; means are shown with stars. Error bars show the standard deviation. (a) The per-participant data reveals considerable individual differences behind the overall accuracy trend (b) Per-color pair data shows that some color pairs were considerably easier than others overall, as well as differences between the effect of condition on accuracy. The pairs are numbered according to their IDs in Figure 3.

### 3.2 Effect of viewing condition on accuracy

We next tested whether these differences between viewing conditions were statistically reliable once differences between participants and between color pairs were taken into account. A generalized linear mixed model was fit to model the effect of viewing condition on binary response accuracy. The model included fixed effects of viewing condition (V) and color pair (C), and a random effect for participant (P):

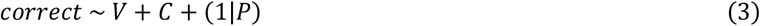

Including color pair significantly improved model fit, *χ*^2^(7) = 44.4, p < 0.001, and reduced AIC from 1267.1 to 1236.7. Compared to free viewing, hard-fixed viewing significantly reduced the odds of making a correct response (*β* = −0.66, *SE* = 0.17, *z* = −3.94, p < 0.001, odds ratio (OR) = 0.52 [0.37, 0.72]). Accuracy under easy-fixed viewing did not differ significantly from free viewing (*β* = 0.17, *SE* = 0.17, *z* = 0.97, p = 0.33, OR = 1.18 [0.84, 1.67]). In other words, being able to tilt the surfaces nearly doubled the odds of a correct response relative to viewing them at an uninformative fixed tilt, and brought performance to a level statistically indistinguishable from viewing them at an informative fixed tilt.

### 3.3 Effect of trajectory summary measures on accuracy

Having established that free tilting helps, we next asked how it helps. The simplest possibility is that performance depends on how much color-difference information observers encountered while tilting. Each of the three summary measures considered below captures a different way in which observers might use the sampled information: the mean Δ*I* reflects the evidence accumulated over the whole trajectory, the peak Δ*I* reflects the maximum evidence encountered at any point, and the end Δ*I* reflects the evidence available at the tilt where the participant was ready to report the decision. For peak Δ*I*, we took the 95th percentile of encountered color difference: a single peak value is noisy and could have lasted only a single IMU sample (0.016 s). We analyzed whether these measures could predict free-viewing accuracy. Each measure was *z*-scored and included as a fixed effect (S) in a separate GLMM alongside fixed effects for trial duration (T) and color pair (C) and a random effect for participant (P) as controls. The three summary measures are correlated and so we avoided including them within the same model.

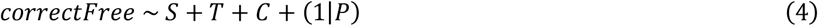

Only mean and end Δ*I* improved model fit relative to a baseline model with control effects (p < 0.05). Model fit comparisons are shown in Table 1.

**Table 1.** Model fit comparison between sampled Δ*I* summary metrics.

| Model | Log-Likelihood | AIC | Δ AIC | χ<sup>2</sup>(1) | p |
|---|---|---|---|---|---|
| Baseline | −295.7 | 611.3 | — | — | — |
| + mean Δ*I* | −293.3 | 608.5 | −2.8 | 4.82 | 0.028 |
| + peak Δ*I* | −294.2 | 610.3 | −1.0 | 2.97 | 0.085 |
| + end Δ*I* | −292.8 | 607.5 | −3.8 | 5.78 | 0.016 |

Mean Δ*I* and end Δ*I* both positively predicted accuracy, whereas peak Δ*I* had no significant effect (parameters shown in Table 2). Duration (M = 19.9s, SD = 12.8s) negatively predicted accuracy (with mean Δ*I*, β = −0.346; p < 0.01; with end Δ*I*, β = −0.342, p < 0.01). The latter effect is likely because harder trials elicited both longer search times and poorer accuracy. Thus, free-viewing accuracy was higher on trials in which observers spent more time at informative tilts and finished at an informative tilt, whereas merely passing through a highly informative tilt at some point did not reliably predict accuracy.

**Table 2.** Fixed effects parameters for sampled Δ*I* summary metrics.

| Parameter | β | SE | *z* | *p* | OR [95% CI] |
|---|---|---|---|---|---|
| Mean Δ*I* | +0.249 | 0.108 | 2.21 | 0.027 | 1.27 [1.03, 1.57] |
| End Δ*I* | +0.256 | 0.106 | 2.43 | 0.016 | 1.29 [1.05, 1.59] |
| Peak Δ*I* | +0.201 | 0.116 | 1.73 | 0.084 (ns) | 1.22 [0.97, 1.54] |

### 3.4 Motion alignment with discriminability gradients

The summary measures above describe how much information observers sampled, but not whether their movements were directed toward informative tilts. In this section we ask whether observers tilted the platform along the directions in which the color difference changes most (Fig. 5a). To do this, we compare the direction of each movement segment with the local gradient of ΔI, and summarize the agreement over a whole trajectory by a single alignment score.

**Figure 5.**
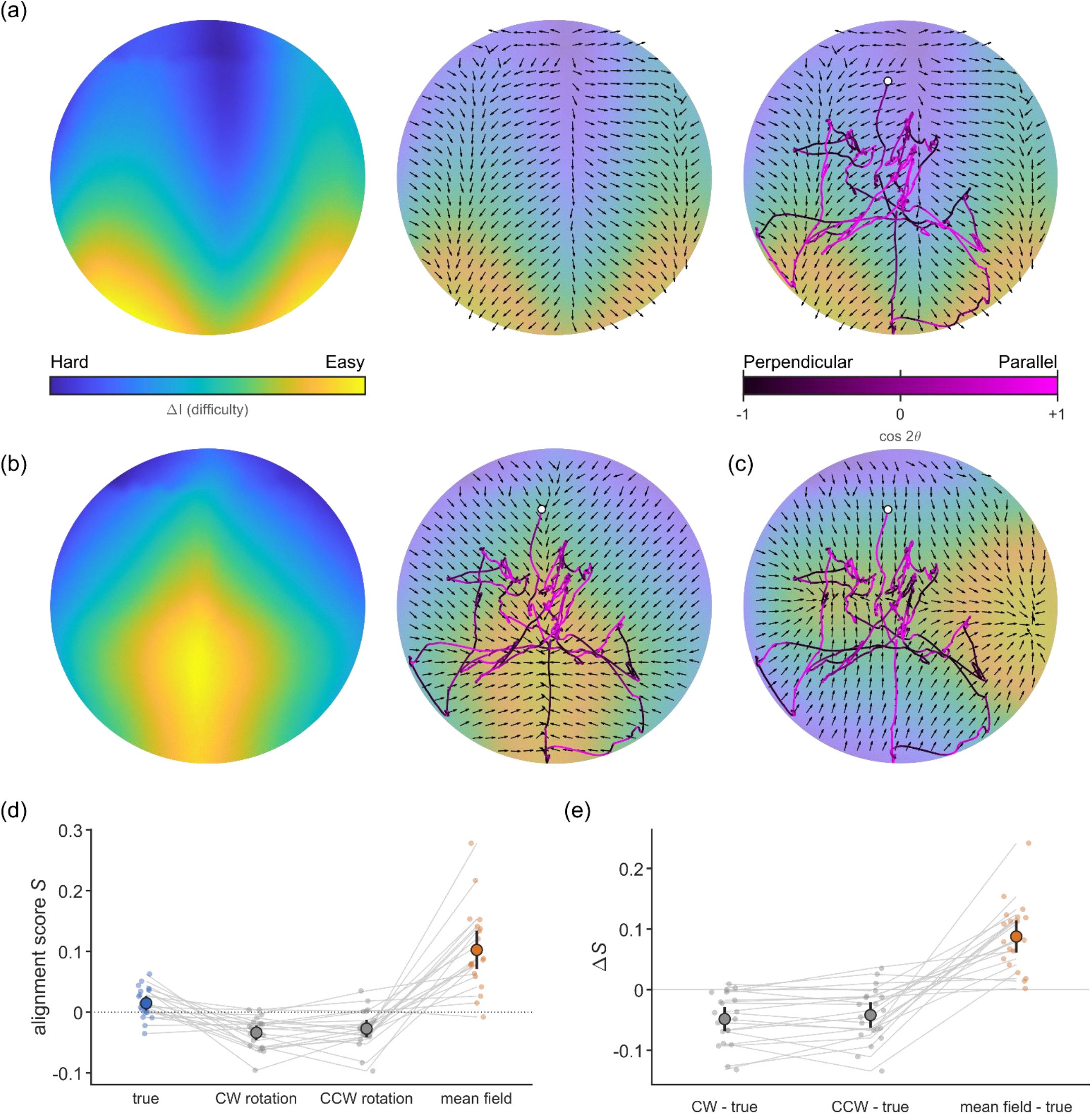
Visualization of the alignment analysis. a) The heatmap on the left shows the spatial distribution of color difference signals at each tilt. The arrows show which direction maximizes color difference (the gradient). The right figure shows an example trajectory, starting at the white dot, and colored according to how well the movement aligns with the color difference gradient. b) The 90° clockwise rotated heatmap, and the same trajectory’s alignment with it. c) The mean of all heatmaps in the warm–cool configuration, and the same trajectory’s alignment with it. The color scales are normalized to the minimum and maximum color difference in panels (a)–(e). d) Each participant’s average alignment score across their trials. Lines connect the scores for the same participant. The error bars show the standard error of the mean. e) Each participant’s average difference score for the three comparisons: clockwise – true; counterclockwise – true; mean – true. The line formatting is the same as for panel (d).

For a given stimulus configuration and lighting orientation, Δ*I* is a scalar field *f*(*Lx, Ly, σ*) primarily dependent on tilt position (*Lx, Ly*) but also changes with spin *σ*. The gradient

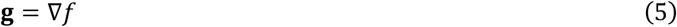

defines the *tilt* direction that maximally increases Δ*I* with spin held constant. For each recorded segment *t*, the alignment between the motion direction **v**_*t*_ and gradient **g**_*t*_ is given by:

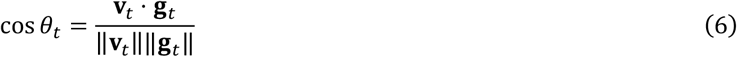

where *θ*_*t*_ is the angle between the movement and gradient vectors for segment *t*. As tilt trajectories exist on a sphere, we used the pullback metric of the sphere to correct for the curvature of the spherical surface.

To quantify the overall alignment between trajectories and their corresponding information fields, we calculated an order parameter [28] for each trajectory:

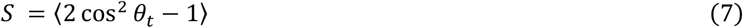

where ⟨ ⋅ ⟩ is the mean over all segments. The order parameter is an alignment score that measures unsigned alignment (i.e.collinearity) with a given direction (the gradient), regardless of the sign of the movement. This prevents the average value collapsing to zero, which would otherwise be observed during oscillatory motion that is well-aligned. A value of +1 means perfect parallel alignment, 0 means random alignment, and −1 means perfect perpendicular alignment. We then averaged the order parameters within each participant, then across all participants.

To test whether the trajectories were aligned with the stimulus-specific distribution of Δ*I*, we compared alignment between the observed Δ*I* field for a given trial and either (i) simulated fields under a rotated lighting axis (90° clockwise and counterclockwise; Fig. 5b); or (ii) the mean Δ*I* field across all 48 stimulus configurations (Fig. 5c). The former tests whether the trajectory is aligned with the specific field in that orientation (i.e. that specific color pair under that specific lighting axis); the latter tests whether trajectories are more aligned to the field than the generic spatial structure of Δ*I* imposed by the lighting axis. Fig. 5d shows each participant’s averaged alignment scores with each field. Alignment with the true and rotated fields were consistent and close to zero. However, alignment with the mean field was generally larger and showed more variation between participants. Some individual participants’ alignment scores were consistently near zero across fields, corresponding to random motion relative to each field. Table 3 shows the average alignment scores across participants for each field, tested against chance. Using the Holm method to control for multiple comparisons, participants’ average alignment was different from zero for all fields.

**Table 3.** Mean trajectory alignment with each field, tested against zero. The 95% confidence intervals (CIs), standard deviations (SDs), and results of two-tailed one-sample t-tests against zero are shown for each field.

| Field | Average alignment score S | 95% CI | SD | *t(19)* | *p* |
|---|---|---|---|---|---|
| True field | +0.015 | [+0.003, +0.027] | 0.025 | +2.62 | 0.02 |
| Clockwise | −0.034 | [−0.046, −0.022] | 0.026 | −5.79 | <0.001 |
| Anticlockwise | −0.027 | [−0.042, −0.013] | 0.031 | −3.91 | 0.002 |
| Mean field | +0.103 | [+0.071, +0.134] | 0.068 | +6.73 | <0.001 |

For each trajectory, we calculated the difference in alignment between the true and alternative fields. We averaged these differences within each participant. Fig. 5e shows each participant’s average alignment differences between fields. For each alternative field, we performed a one-sample *t*-test to assess whether there was a difference between alignment to the true field and the alternative field. Table 4 shows the results of alignment comparison. Alignment with the actual field was greater than with the rotated fields. However, trajectories were more aligned with the mean field across stimulus configurations than each trial’s actual field. In other words, observers did tend to sample axes that are informative on average under a given lighting arrangement, but there was little evidence that they tailored their movements to the specific color pair presented on a given trial.

**Table 4.** Alignment differences between each field and the true field.

| Field | Mean difference (true minus alternate) | 95% CI | SD | *t(19)* | *p* |
|---|---|---|---|---|---|
| Clockwise | +0.048 | [+0.029, +0.068] | 0.043 | +5.08 | <0.001 |
| Anticlockwise | +0.042 | [+0.021, +0.063] | 0.045 | +4.16 | <0.001 |
| Mean field | −0.088 | [−0.114, −0.061] | 0.057 | −6.93 | <0.001 |

### 3.5 Frequency analysis

Here we ask whether observers’ sampling had a temporal structure, i.e. whether the platform was moved in a rhythmic, repetitive way (for example, moving back and forth between two tilts) rather than moving randomly, and if so, whether this rhythm was also present in the sampled color-difference signal. The logic is that unstructured movement (such as Brownian motion) is expected to produce a power spectrum that falls smoothly with frequency (a 1/*f*^2^ spectrum) [29], whereas systematic repetitive motion adds power at particular frequencies, which appears as a deviation from this baseline.

Thus, we analyzed the spectral features of three variables: *Lx* and *Ly* motion and the sampled Δ*I*. For each trial, we computed the discrete Fourier transform and estimated the one-sided power spectral density (PSD) *P*(*f*_*k*_) as:

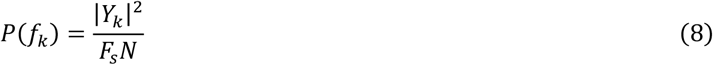

where *Y*_*k*_ is the Fourier component at frequency *f*_*k*_, *F*_*s*_ is the IMU sampling frequency (62.5 Hz) and *N* is the number of samples in each trial. To compare power spectra across variable-length trials, we estimated the PSD in 0.15 Hz-wide frequency bands, whose centers ranged from 0.10 Hz to 31.15 Hz (see Fig. 6a). For each frequency band *B*_*b*_, the mean PSD was calculated from the Fourier frequencies falling within the band and assigned to the band’s center frequency *f*_*b*_ as:

**Figure 6.**
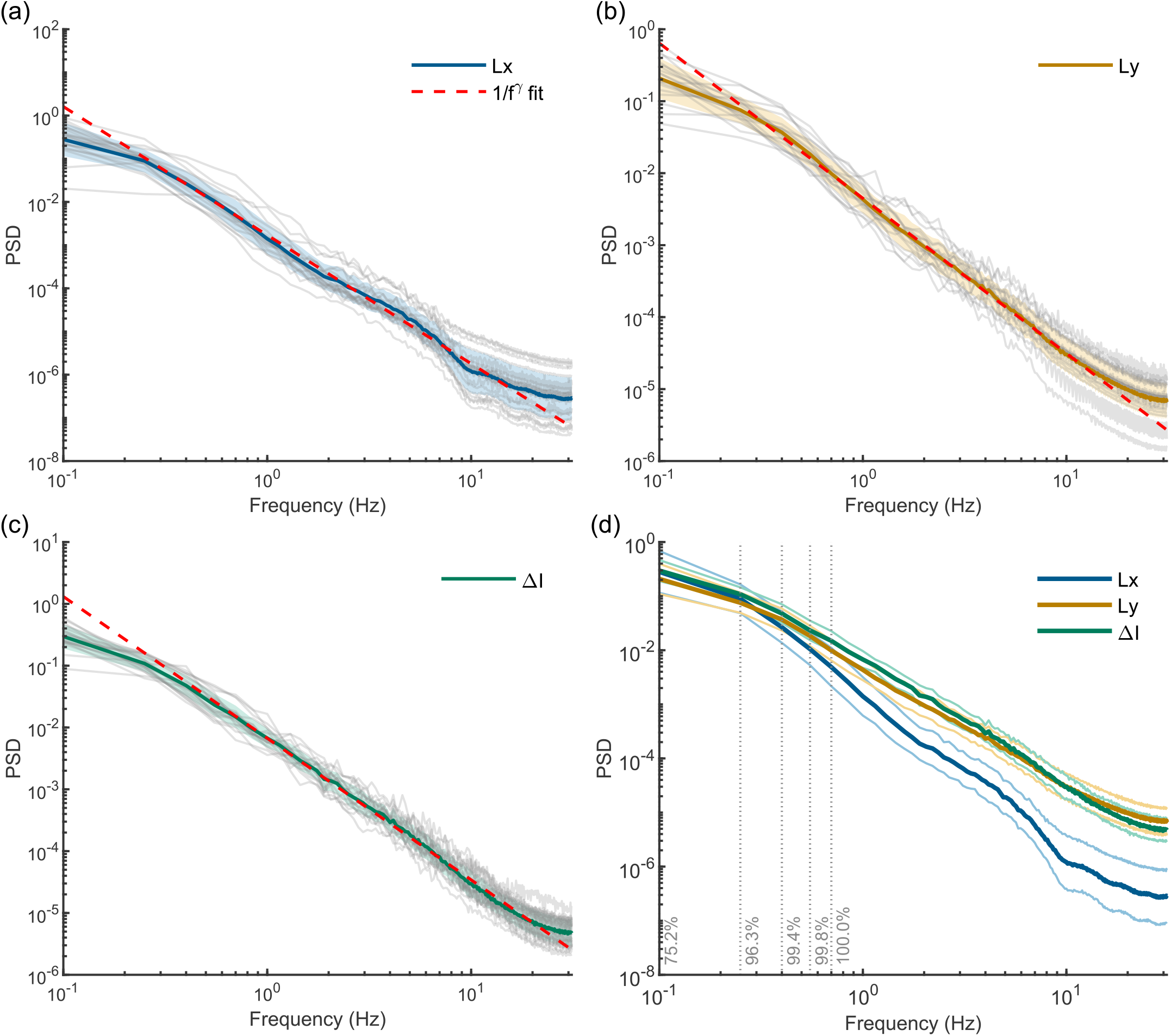
(a) Mean power spectra across participants. The solid line shows the mean of participants’ power spectra. The shaded bands show the standard deviation of estimated PSD. The dashed line is the average 1/*f*^*γ*^ fit for each variable. The labeled dotted lines indicate the proportion of trials lasting at least one cycle at that frequency, i.e. contributed to that PSD estimate. (b) The mean power spectrum for each variable is shown independently. The light gray lines are participants’ power spectra.

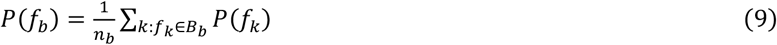

The result is three power spectra for each trial, estimated over a shared range of frequencies for *Lx*(t), *Ly*(t), and Δ*I*(t). Power spectra were averaged within each participant. Fig. 6a shows the mean participant power spectra across participants for each variable. These spectra were then fitted with a power-law model:

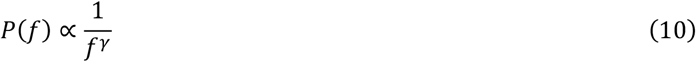

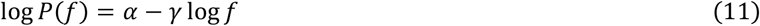

We used linear regression to estimate *α* and *γ* over a narrower frequency window of 0.25 Hz to 10 Hz. The physical constraints of the tripod head’s range of motion prevent tilt position. Therefore, deviations from a Brownian 1/*f*^2^ scaling at lower frequencies are expected [29, 30]; a process that is bounded might require a different stochastic model for very low frequencies [31]. We assumed that higher frequencies primarily reflected physiological, biomechanical or measurement related processes, rather than meaningful manipulations of the platform. The exponent *γ* describes the rate at which power decreases with increasing frequency. These were modeled for each participant’s average power spectrum. For each variable, the average exponents across participants were as follows: *Lx(t), γ* = 2.98, *SD* = 0.32; *Ly*(*t*), *γ* = 2.15, *SD* = 0.14; *ΔI*(*t*), *γ* = 2.29, *SD* = 0.12. *Lx* showed a greater decrease in power at higher frequencies, suggesting slightly more structured motion compared to a random baseline, while *Ly* and *ΔI* resembled a random walk more closely.

## 4. Discussion

### 4.1 The present findings

This experiment demonstrates that human observers can utilize surface tilt to improve surface color discrimination. Surfaces that are metameric under one illumination to an observer with trichromatic human color vision may be disambiguated under a second illumination. Real environments have directional spectral variation that can feasibly break metamerism in this way if observers are able to sample spectrally differing illuminations through surface tilt [5,7]. This experiment demonstrates that human observers can utilize surface tilt to improve surface color discrimination. The results are consistent with Morimoto et al.’s [6] simulation: compared to a relatively indiscriminable baseline, color discrimination significantly improved when participants were able to tilt surfaces, and performance became comparable to viewing relatively discriminable surfaces.

An additional question was how color difference signals were sampled through surface tilt. One hypothetical extreme is that participants would explore randomly and answer as soon as they reach a suprathreshold tilt. Alternatively, they may integrate evidence accumulated over a whole random trajectory and answer accordingly. A third possibility is that observers’ motions are directed toward tilts that specifically help disambiguate colors. The behaviors associated with these strategies would be (a) responding to incidental informative evidence, (b) responding based on exposure history and (c) intentionally moving toward more informative tilts. The final strategy—though idealized—would be optimal, but requires the observer to anticipate the spatial distribution of color signals for a given sample configuration under the lighting environment.

The peak color difference encountered had no significant effect on performance. This does not necessarily rule out the possibility that participants respond based on positively skewed evidence. For example, the greatest signals may not have been integrated if they were encountered in transit between different dwells. They may have been difficult to find or perceive, such as if they occurred at grazing or extreme tilts. The color difference at the end tilt did predict improved performance. The most plausible explanation is that participants are more likely to stop sampling and make a correct decision at a tilt where the surfaces are distinct. However, as some participants would continue to move the platform while answering, such as resetting the platform to a default tilt, or placing it out-of-view to prepare for the next trial, or continuing to sample tilts, the end-of-trajectory data may not necessarily reflect the final sampled color difference. Mean encountered color difference also predicted correct responses and was a more robust, simple summary of the sampled color difference signals. These findings suggest that being able to sample more informative tilts does improve color discrimination, but not just by chancing upon tilts with incidentally higher discriminability.

Trajectories were most aligned with the average distribution of color signals across all surface configurations. The corresponding alignment score of +0.103 suggests a subtle preference toward sampling the directions that generally change surface color appearance under the lighting environment. This effect could also be due to learning which tilts tend to discriminate surfaces under a given lighting arrangement. Participants were only slightly aligned with the stimulus-specific distribution of color information compared to chance. It is unfeasible for participants to intuit a priori which tilts are most informative for a given stimulus. The difficulty of searching optimally on each trial may have been exacerbated by the overall difficulty of the task, which was chosen to be near threshold. Participants would instead prefer to sample the axes that are generally discriminating for a range of surfaces; in our experiment, these were the tilts that faced each light source directly (Fig. 5c).

The frequency analysis suggested that movement in the left–right direction had proportionally more power in lower frequencies compared to a random walk, whereas movement in the front–back direction, as well as in the sequence of sampled Δ*I*, appeared closer to a random walk. This would corroborate the idea that participants may manipulate the platform systematically with respect to the general lighting geometry, since the left–right axis aligns with the principal axis of variation in lighting. However, there are also directional asymmetries in the physical constraints, which may affect the observed power. The practical range of the front–back axis is relatively narrow as many tilts are out of view. Moreover, left–right and front– back motion require different physical motor demands. Although power spectra alone cannot definitively identify the underlying stochastic dynamics, they may suggest that there is an asymmetry and temporal structure in the movement dynamics. The findings of both trajectory analyses warrant a future experiment with a different lighting arrangement, which would empirically test whether participants systematically sample the general lighting structure, and to test whether any such tendency is purely a motion preference.

### 4.2 Implications

The number of information channels in human color vision is limited by the number of cone classes. Given a surface’s tristimulus values under one illumination, it would be impossible to predict its complete reflectance properties [20]. As the theoretical extent of this ambiguity is considerable [2], observers could seek to overcome it by accessing more information than present in a single view. Multiple views of a surface—under different illuminations or at different tilts—provide a richer set of cone signals associated with a surface [32]. This variation may provide additional information about the surface reflectance properties, as the cone signals depend jointly on the surface reflectance and the illumination. This idea relates closely to the problem of color constancy: the relatively constant representation of surface color despite changes in illumination. It has been proposed that object-color perception under spatially varying illumination involves additional ‘lighting’ dimensions of the surface color representation beyond the conventional dimensions of trichromatic color perception [2, 33]. Moreover, different surface reflectances vary systematically in how their color appearance changes with illumination [34]. Tilting a surface may therefore allow observers to sample different illumination conditions and obtain information that cannot be ascertained from a single view. The finding that observers preferentially sample axes that produce larger changes in color appearance may reflect sensitivity to such informative variations.

Human observers readily exploit other sources of information beyond ratios of fixed-sensitivity cone cells during color perception. Dichromats, for example, may exploit rod responses, lightness cues, or differences in spectral sensitivities for different stimulus sizes [35, 36]. In practice, functional color vision is additionally influenced by cues such as 3D shape [37– 39], specular highlights [40], and, over longer time periods, color memory [41, 42]. Several studies have shown that observers make better judgments about material properties under dynamic viewing [43–45]. Increasingly, evidence suggests that observers can intentionally sample transformations that provide property-disambiguating information, and that some of their strategies are dependent on the current task [14, 15]. Very little empirical evidence exists regarding dynamic cues and active sensing in color vision. This study provides an empirical account of how humans can actively improve color discrimination through surface tilt: one potential strategy is sampling the direction of maximum illuminant variation.

### 4.3 Limitations and extensions

The present study is not free from limitations. The tripod’s range of motion mechanically constrains the available tilts to the upper hemisphere. Across this range of motion, both lights are always illuminating the surface with minimal interreflection. This structure differs considerably from measured environmental lighting, where upper and lower hemisphere illumination can vary dramatically [5]. The information metric is also not perfect. Due to the geometry of the setup, our simulated Δ*I* can occasionally take on slightly negative values, corresponding to a distractor appearing more distinct than the target patch. The Δ*I* represents how much evidence there is for the correct patch; any analysis of Δ*I* ignores situations where there is more evidence for the distractor patches, which could be an explanation for poorer performance in free-viewing. Any tilting in our setup continuously resamples different proportions of the two direct illuminants, whereas a less constrained environment might have more discrete spectral differences between illuminants sampled at adjacent tilts; observers can more dramatically change the incident illumination by shading a surface, such as by tilting it completely away from, or occluding it from a direct illuminant. Nevertheless, the results imply that, even in the minimal case where only continuous illuminant differences can be sampled, observers still benefit from active tilting. In addition to the spatially idealized configuration, the experiment only used a warm–cool axis with a limited selection of surface pairs, which would affect whether the mean heatmap finding is generalizable to all possible surfaces. Color discrimination is poorer under environmental illumination with atypical chromaticities, such as magenta-green combinations [46]. Finally, not all objects are easily manipulable, such as large or distant objects; in such cases, natural changes in lighting geometry, such as occlusion by clouds, moving shadows, or moving light sources may provide an alternative means of sampling informative variations in visual appearance. More complex surfaces might also be disambiguated by changes in viewpoint. The results from our lab experiment may not fully generalize to the diversity of surfaces, objects and illuminants observers might encounter.

### 4.4 Summary

Ultimately, the study shows that human observers can improve color discrimination by tilting surfaces under directional illumination. Participants appeared to sample the general lighting structure rather than the exact distribution of color signals for a given surface configuration. The results complement anecdotal [7] and simulated [6] evidence suggesting that surface tilt is a feasible and accessible way of actively improving color discrimination. The results also highlight yet again the importance of ‘natural environments’ in color vision research—in our case, directionally varying illumination where observers can freely tilt surfaces.

## Acknowledgment

The authors would like to acknowledge Dr David Brainard for suggesting the separation between easy and hard viewing conditions. SM and FC’s posts were supported by the Anne Silk Legacy (awarded to HES). This work was supported by a Wellcome Trust Research Enrichment – Diversity & Inclusion Grant (218657/Z/19/A) awarded to TM.

## Funding

### Disclosures

The authors declare no conflicts of interest.

### Data availability

The data are available from the authors upon reasonable request.

## Notes

### Competing Interest Statement

The authors have declared no competing interest.

## References

1. W. A. Rushton, “Pigments and signals in colour vision,” J. Physiol. 220, 1P–31P (1972). 10.1113/jphysiol.1972.sp009719

2. A. D. Logvinenko, B. Funt, H. Mirzaei, and R. Tokunaga, “Rethinking colour constancy,” PLoS ONE 10, e0135029 (2015). 10.1371/journal.pone.0135029

3. D. H. Foster, K. Amano, S. M. C. Nascimento, and M. J. Foster, “Frequency of metamerism in natural scenes,” J. Opt. Soc. Am. A 23, 2359–2372 (2006).

4. A. Akbarinia and K. R. Gegenfurtner, “Color metamerism and the structure of illuminant space,” J. Opt. Soc. Am. A 35, B231–B238 (2018).

5. T. Morimoto, S. Kishigami, J. M. M. Linhares, et al., “Hyperspectral environmental illumination maps: characterizing directional spectral variation in natural environments,” Opt. Express 27, 32277–32293 (2019).

6. T. Morimoto, J. M. M. Linhares, S. M. C. Nascimento, and H. E. Smithson, “How many surfaces can you distinguish by color? Real environmental lighting increases discriminability of surface colors,” Opt. Express 32, 34246–34253 (2024).

7. J. Broackes, “What do the color-blind see?” in Color Ontology and Color Science (MIT Press, 2010), pp. 291–405.

8. J. J. Gibson, The Perception of the Visual World (Greenwood, 1974).

9. J. J. Gibson, The Senses Considered as Perceptual Systems (Greenwood, 1983).

10. D. H. Foster, K. Amano, and S. M. C. Nascimento, “Colour constancy from temporal cues: better matches with less variability under fast illuminant changes,” Vision Res. 41, 285–293 (2001).

11. M. D’Zmura and A. Mangalick, “Detection of contrary chromatic change,” J. Opt. Soc. Am. A 11, 543–546 (1994).

12. A. Gauch and D. Kerzel, “Perceptual asynchronies between color and motion at the onset of motion and along the motion trajectory,” Percept. Psychophys. 70, 1092–1103 (2008).

13. B. Xiao and C. Liao, “Material perception connects vision, cognition and action,” Nat. Rev. Psychol. 4, 687–701 (2025).

14. R. Nomachi, H. Tamura, K. A. Helgeland, et al., “Material discrimination relies on context-dependent active sensing strategies,” J. Vis. 26(6), 4 (2026).

15. L. P. Y. Lin, K. Drewing, and K. Doerschner, “Exploring for gloss: active exploration in visual material perception,” J. Vis. 25(6), 2 (2025).

16. M. Rucci and J. D. Victor, “The unsteady eye: an information-processing stage, not a bug,” Trends Neurosci. 38, 195–206 (2015).

17. J. M. Findlay and I. D. Gilchrist, Active Vision: The Psychology of Looking and Seeing (Oxford University Press, 2003).

18. M. Land and B. Tatler, Looking and Acting: Vision and Eye Movements in Natural Behaviour (Oxford University Press, 2009).

19. K. Friston, T. FitzGerald, F. Rigoli, et al., “Active inference and learning,” Neurosci. Biobehav. Rev. 68, 862–879 (2016).

20. L. Hardy, G. Rand, and M. C. Rittler, HRR Pseudoisochromatic Plates, 4th ed. (Richmond Products, 2002).

21. W. Jakob, S. Speierer, N. Roussel, et al., “Mitsuba 3 renderer,” version 3.7.1, https://mitsuba-renderer.org.

22. G. Wyszecki and W. S. Stiles, Color Science: Concepts and Methods, Quantitative Data and Formulae, 2nd ed. (Wiley, 1982).

23. International Color Consortium, Specification ICC.1:2004-10 (Profile Version 4.2.0.0): Image Technology Colour Management—Architecture, Profile Format and Data Structure (2006). https://www.color.org/icc1v42.pdf

24. The MathWorks, Inc., MATLAB, version 24.2.0.2923080 (2024).

25. M. Kleiner, D. Brainard, D. Pelli, et al., “What’s new in Psychtoolbox-3,” Perception 36, 1–16 (2007).

26. G. Sharma, W. Wu, and E. N. Dalal, “The CIEDE2000 color-difference formula: implementation notes, supplementary test data, and mathematical observations,” Color Res. Appl. 30, 21–30 (2005).

27. L. Arend and A. Reeves, “Simultaneous color constancy,” J. Opt. Soc. Am. A 3, 1743–1751 (1986).

28. A. A. Mercurieva and T. M. Birshtein, “Liquid-crystalline ordering in two-dimensional systems with discrete symmetry,” Macromol. Theory Simul. 1, 205–214 (1992).

29. F. Tessari, J. Hermus, R. Sugimoto-Dimitrova, et al., “Brownian processes in human motor control support descending neural velocity commands,” Sci. Rep. 14, 8341 (2024).

30. J. J. Collins and C. J. De Luca, “Open-loop and closed-loop control of posture: a random-walk analysis of center-of-pressure trajectories,” Exp. Brain Res. 95, 308–318 (1993). 10.1007/BF00229788

31. J. L. Doob, “The Brownian movement and stochastic equations,” Ann. Math. 43, 351–369 (1942).

32. M. D’Zmura and A. Mangalick, “Detection of contrary chromatic change,” J. Opt. Soc. Am. A 11, 543–546 (1994). 10.1364/JOSAA.11.000543

33. R. Tokunaga and A. D. Logvinenko, “Material and lighting dimensions of object colour,” Vision Res. 50, 1740–1747 (2010).

34. D. L. Philipona and J. K. O’Regan, “Color naming, unique hues, and hue cancellation predicted from singularities in reflection properties,” Vis. Neurosci. 23, 331–339 (2006).

35. K. Uchikawa, “Trichromat-like categorical color naming of dichromats,” VISION 20, 62–66 (2008).

36. E. D. Montag and R. M. Boynton, “Rod influence in dichromatic surface color perception,” Vision Res. 27, 2153–2162 (1987).

37. M. G. Bloj, D. Kersten, and A. C. Hurlbert, “Perception of three-dimensional shape influences colour perception through mutual illumination,” Nature 402, 877–879 (1999).

38. M. Hedrich, M. Bloj, and A. I. Ruppertsberg, “Color constancy improves for real 3D objects,” J. Vis. 9(4), 16 (2009).

39. B. Xiao, B. Hurst, L. MacIntyre, and D. H. Brainard, “The color constancy of three-dimensional objects,” J. Vis. 12(4), 6 (2012).

40. T. Nagai, S. Kaneko, Y. Kawashima, and Y. Yamauchi, “Do specular highlights and the daylight locus act as cues for estimating illumination color from a single object?” Opt. Rev. 24, 47–61 (2017).

41. K. R. Gegenfurtner, D. Weiss, and M. Bloj, “Color constancy in real-world settings,” J. Vis. 24(2), 12 (2024).

42. M. Olkkonen, T. Hansen, and K. R. Gegenfurtner, “Color appearance of familiar objects: effects of object shape, texture, and illumination changes,” J. Vis. 8(5), 13 (2008).

43. T. Young, “The Bakerian lecture: on the theory of light and colours,” Philos. Trans. R. Soc. Lond. 92, 12–48 (1802).

44. K. Doerschner, R. W. Fleming, O. Yilmaz, et al., “Visual motion and the perception of surface material,” Curr. Biol. 21, 2010–2016 (2011).

45. A. Malik, Y. Yu, H. Boyaci, and K. Doerschner, “Perceiving material qualities from moving contours,” bioRxiv (2025). 10.1101/2025.07.21.665763

46. J. J. R. van Assen, P. Barla, and R. W. Fleming, “Visual features in the perception of liquids,” Curr. Biol. 28, 452–458.e4 (2018)

47. T. Morimoto and H. E. Smithson, “Discrimination of spectral reflectance under environmental illumination,” J. Opt. Soc. Am. A 35, B244–B255 (2018).

